# Genetic heterogeneity in *Anopheles darlingi* related to biting behavior in western Amazon

**DOI:** 10.1101/358556

**Authors:** Melina Campos, Diego Peres Alonso, Jan E. Conn, Joseph M. Vinetz, Kevin J. Emerson, Paulo Eduardo Martins Ribolla

## Abstract

In the Amazon Basin, *Anopheles* (*Nyssorhynchus*) *darlingi* is the most aggressive and effective malaria vector. In endemic areas, behavioral aspects of anopheline species such as host preference, biting time and resting location after a blood meal have a key impact on malaria transmission dynamics and transmission control strategies. *An. darlingi* present a variety in behavior throughout its broad distribution including blood feeding related. To investigate the genetic basis of its biting behaviors, host-seeking *An. darlingi* were collected in two settlements (Granada and Remansinho) in Acre, Brazil. Mosquitoes were classified by captured location (indoors or outdoors) and time (dusk or dawn). Genome-wide SNPs were used to assess the degree of genetic diversity and structure in these groups. There was evidence of genetic component of biting behavior regarding both location and time in this species. This study supports that *An. darlingi* blood-feeding behavior has a genetic component. Additional ecological and genomic studies may help to understand the genetic basis of mosquito behavior and address appropriate surveillance and vector control.

**Author Summary:** Malaria is a disease caused by parasite of the genus *Plasmodium* and is transmitted by mosquitoes of the genus *Anopheles*. In the Amazon Basin, the main malaria vector is *Anopheles darlingi*, which is present in high densities in this region. Egg development requires that females of this mosquito seek hosts for blood meals. *Anopheles* females blood feeding may occur indoor or outdoor the houses and typically from the sunset to dawn. *Anopheles darlingi* in particular present great variability regarding its behaviour, presenting variety of peak biting times and patterns. This work shows that there is a genetic component that partially explains these two behaviors: location of the blood meal (inside or outside the houses) and time of feeding. Single nucleotide polymorphisms (SNPs) scattered throughout the genome of *Anopheles darlingi* showed genetic diversity and structure in these groups. A comprehensive understanding of the genetic basis for mosquito behaviour may support innovative vector surveillance and control strategies.

## Introduction

*Anopheles (Nyssorhynchus) darlingi* is the main neotropical malaria vector in the Amazon Basin due to its role in human *Plasmodium* transmission [1,2]. Following a continued decrease in the number of malaria cases in America from 2005 to 2014, an increase was observed in the last three years in which Brazil was the major contributor [3,4]. In Brazil, transmission remains entrenched in the Amazon Basin, which accounts for 99.5% of the country’s malaria burden [5]. Human migration to this region over the past century has been accompanied by dramatic environmental modifications and the promotion of malaria transmission [6–8]. Deforestation and anthropogenic land changes are known to be associated with increases in *An. darlingi* presence and abundance and thus malaria risk [9–11].

Root (1926) first described *An. darlingi* based on morphological characters of the egg, fourth-instar larva, pupa, male and female adults. Thenceforth, successive studies reported morphological, behavioral, ecological and genetic heterogeneity throughout its broad distribution from Central to South America [12–15]. Despite of that, *An. darlingi* was considered as monophyletic species until Emerson and collaborators [16] proposed the occurrence of three putative species based on well-supported genetic clusters. In that study, Neotropical biogeographical events explained dispersal of *An. darlingi* population, in a continental scale, leading to diversification within this species. However, additional genetic diversity in a minor scale has been shown for this vector by ecological differences [17,18] and seasonality [19].

With regard to mosquito behavior, two aspects are important to determine pathogens transmission to human: anthropophily and endophily. The first is the predilection of the vector to blood-feed on humans instead of other animals and, the second is its preference for living close to human habitats. *An. darlingi* has presented persistently high anthropophily in many areas but substantial variation has been reported in behavior activity regarding endophily and endophagy i.e. blood-feed indoors [20,21]. This species has successfully colonized anthropogenic habitats such as fish tanks, ditches, micro dams and drains besides being found in natural breeding sites such as lakes, rivers and streams [10,22]. Depending on the region, *An. darlingi* was considered essentially exophagic/endophagic or both. It also has presented resting behavior indoors (endophily) or outdoors (exophily) and a wide range of peak of biting times and patterns [23–26].

The drivers of behavioral shifting have been difficult to isolate and pinpoint in anophelines, although a general consensus is that the responses are mainly due to the distribution of long-lasting insecticidal nets (LLINs) and indoor residual spraying (IRS) [27–29]. Vector behavioral heterogeneity could be resulted of genetically differentiated subpopulations or behavioral plasticity i.e. individuals with same genotype adopt different behavior. The present article investigated genetic population diversity in *An. darlingi* at a micro-scale considering biting behavior. More specifically, we explored the genetic basis of this species by blood-feeding location (indoors or outdoors) and time (dusk or dawn) using genome-wide SNPs. Understanding the genetic contribution to mosquito biting behaviors could lead to the development of molecular methods towards vector surveillance and malaria control.

## Results

### SNP genotyping

A total of 128 individual mosquitoes were sequenced from both settlements, Granada and Remansinho. We considered four collection categories: indoor collection at dusk i.e., 18-21 h; indoors at dawn i.e., 03-06 h; outdoors at dusk and outdoors at dawn (Table 1). From 124,841,110 ddRAD tag sequences (NCBI SRA BioProject PRJNA298241), ~104 million sequences passed several levels of quality filtering and 34.9% (+/-9.8 SD) of this set of reads was aligned to the *An. darlingi* genome [32]. An average of 10,107 (+/-4,123 SD) ddRAD loci were genotyped per sample, and 25% presented SNPs (S1 Table). Analyses were conducted in two parts, considering locality of collection (Granada/Remansinho) and place (indoor/outdoor); and time (dusk/dawn) of collection. First, analyses included indoor and outdoor specimens collected at dusk from both settlements, whereas the second analyses included dusk and dawn collection only from Granada. Analysis presented a number of SNPs genotyped in at least 50% of individuals in that set without previous population assignment.

**Table 1.** Number of *Anopheles darlingi* genotyped by settlement, location and time.

| Location | Region | State | Latitude | Longitude | Dusk | Outdoor | Indoor |  |
| --- | --- | --- | --- | --- | --- | --- | --- | --- |
|  |  |  |  |  |  | Dawn | Dusk | Dawn |
| Granada | Acrelandia | Acre | -9.752 | -67.071 | 24 | 30 | 19 | 23 |
| Remansinho | Labrea | Amazon | -9.497 | -66.582 | 16 | - | 16 | - |

### Heterogeneity genetic of *Anopheles darlingi* by biting behavior Location: Indoor vs. Outdoor

Dataset analyses were conducted for indoor and outdoor samples collected in the dusk in Granada and Remansinho. After filtering, 944 SNPs were found in at least 50% of all individuals from the four groups. Principal component analysis (PCA) showed that indoor specimens were concisely grouped together regardless of the collection location, whereas outdoor individuals were found in two different areas in the plot (Fig 2A). Discriminant analysis of principal component (DAPC) showed *K* = 3 as the best number of genetic *clusters* i.e., the lowest Bayesian Information Criterion (BIC) (Fig 2C). Cluster “1” had only outdoor *An. darlingi* specimens from Granada and cluster “2” had nearly all outdoor individuals from Remansinho, whereas “3” included indoor individuals from both locations (Fig 2B). Indoor groups showed a higher percentage of polymorphism compared with outdoor ones (Table 2). Interestingly, the highest pairwise *FST* was between indoor Remansinho and outdoor Granada samples (*F_ST_* = 0.177, p < 0.0001), and the lowest was between indoors samples (*F_ST_* = 0.094, p <0.0001) (Table 4).

**Figure 1.**
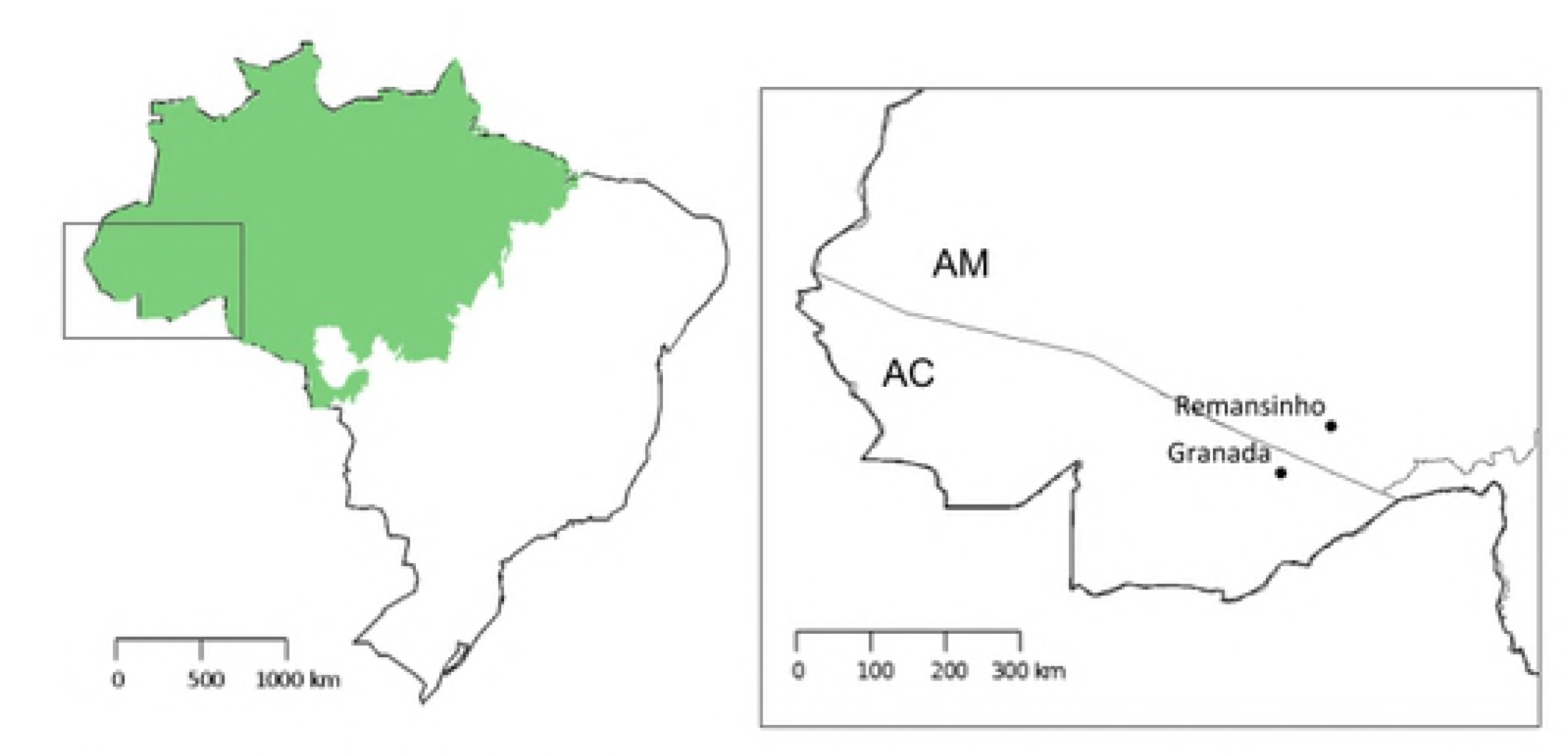
Collection points in Brazilian Amazon Basin. Map of Brazil showing Amazon domain. Square box: zoom in showing Acre (AC) and Amazonas (AM) states and the two collection localities, Granada and Remansinho.

**Figure 2.**
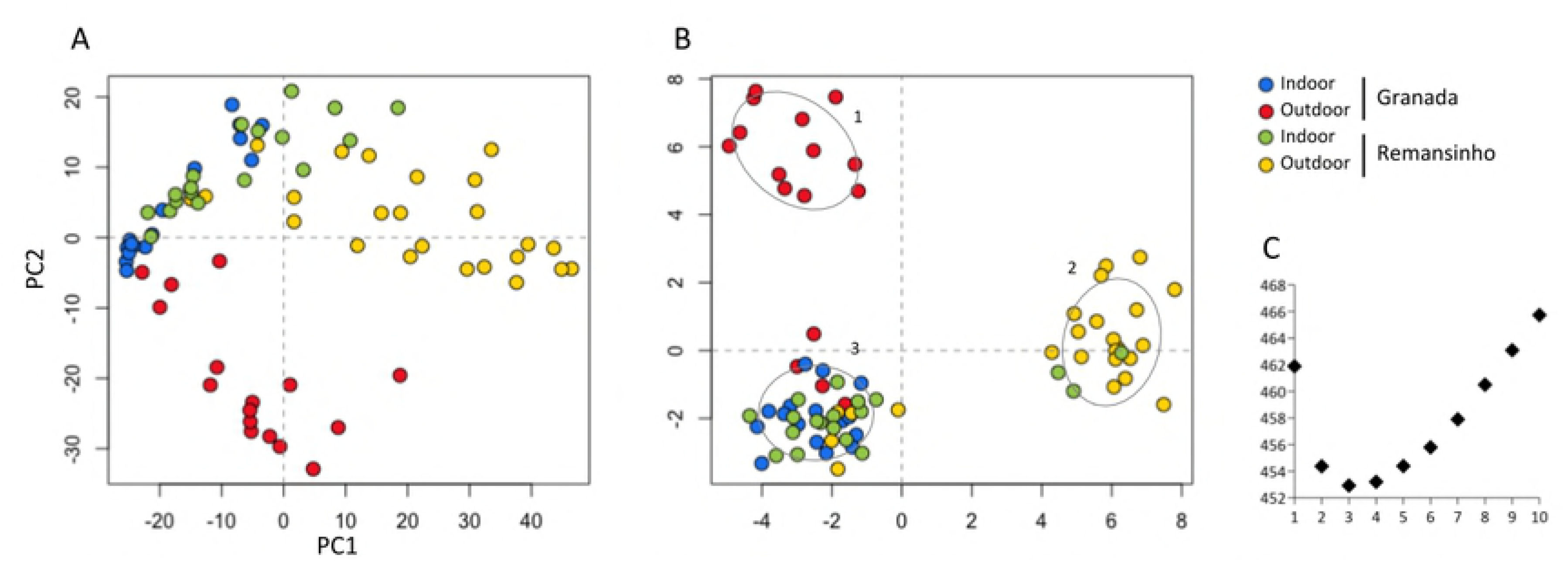
Principal Component Analysis (PCA) and Discriminant analysis of principal components (DAPC) of outdoor and indoor *An. darlingi* genotypes. (A) PCA of outdoor samples (red and yellow dots) and indoor samples (blue and green dots) from Granada and Remansinho. (B) DAPC - ordination of all samples in the three genetic clusters (1-3) in two axes. (C) Bayesian Information Criterion (BIC) to estimate the appropriate number of genetic clusters, which is the lowest BIC value.

**Table 2.**
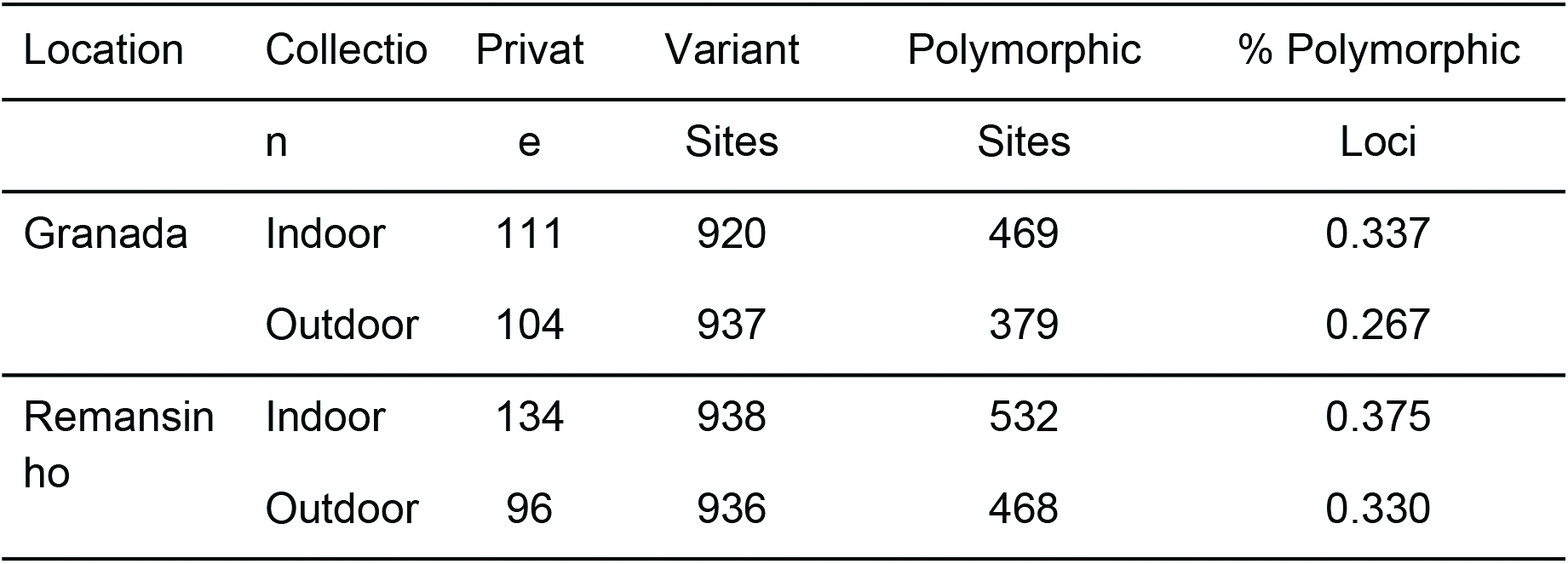
Summary statistics for the indoor and outdoor *Anopheles darlingi*.

### Time: dusk vs. dawn

Four groups of samples, indoors and outdoors collected in the dusk and dawn, were analyzed in this set. A total of 589 SNPs was filtered in at least 50% in dusk/dawn and indoor/outdoor samples. The first axis in the PCA divided indoor and outdoor samples, whereas the second axis separated time of collection (Fig 3A). The lowest BIC for *K*-means was 4 in the DAPC where essentially the four groups were partitioned (Fig 3B-C). Samples collected in the dawn showed a higher percentage of polymorphic loci compared with dusk ones (Table 3). For this dataset, the highest pairwise *F_ST_* was between outdoor dusk and indoor dawn samples (*F_ST_* = 0.259, p < 0.0001), and the lowest was between outdoors samples (*F_ST_* = 0.081, p <0.0001) (Table 5).

**Figure 3.**
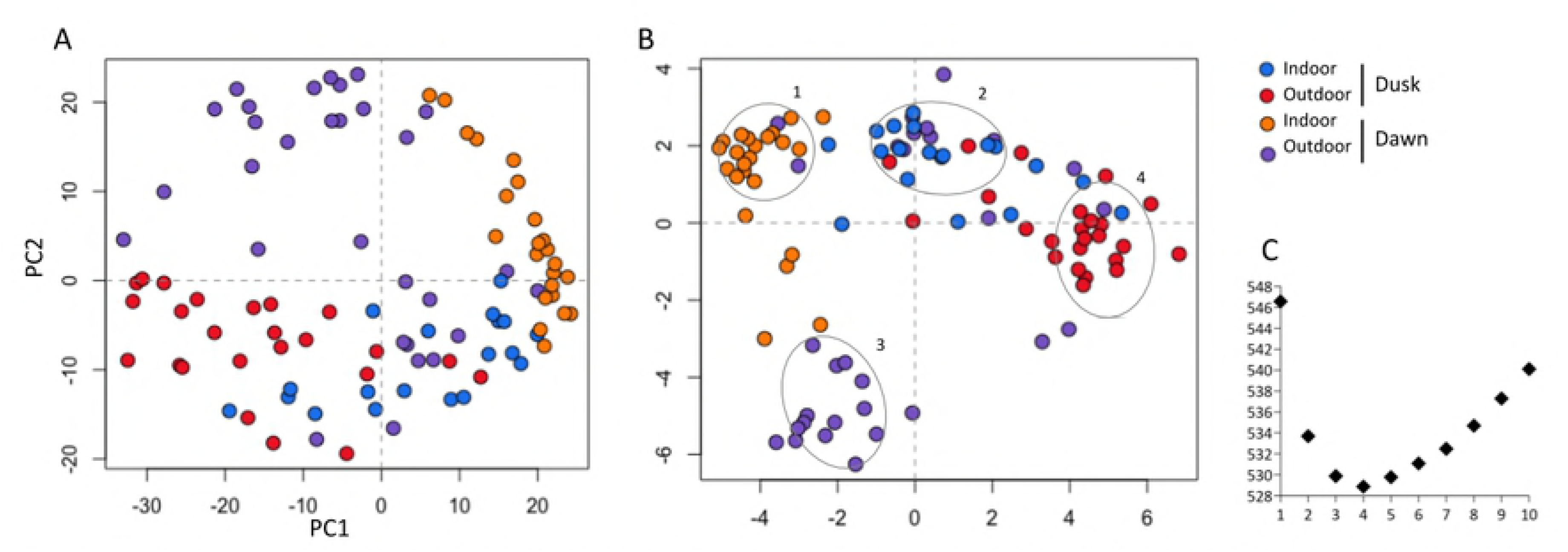
Principal Component Analysis (PCA) and Discriminant analysis of principal components (DAPC) of outdoor/indoor and dusk/dawn *An. darlingi* genotypes. (A) PCA of dusk samples (red and blue dots) and dawn samples (purple and orange dots) from Granada and Remansinho. (B) DAPC - ordination of all samples in the four genetic clusters (1-4) in two axes. (C) Bayesian Information Criterion (BIC) to estimate the appropriate number of genetic clusters, which is the lowest BIC value.

**Table 3.** Summary statistics for the indoor/outdoor and dusk/dawn *Anopheles darlingi*.

| Location | Time | Private | Variant Sites | Polymorphic Sites | % Polymorphic Loci |
| --- | --- | --- | --- | --- | --- |
| Indoor | Dusk | 55 | 556 | 243 | 0.289 |
|  | Dawn | 85 | 557 | 315 | 0.374 |
| Outdoor | Dusk | 70 | 556 | 190 | 0.226 |
|  | Dawn | 91 | 557 | 291 | 0.346 |

**Table 4.**
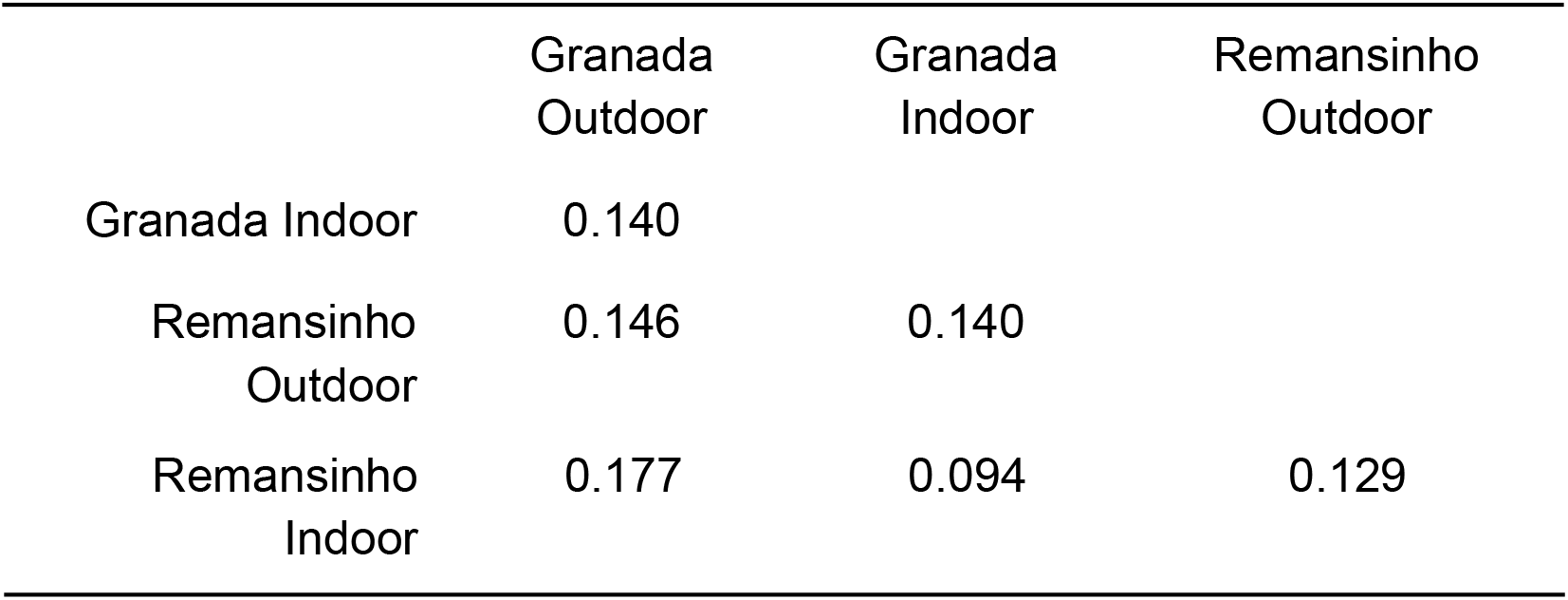
Pairwise *F_ST_* between indoor and outdoor *Anopheles darling* i.

**Table 5.** Pairwise *F_ST_* between indoor/outdoor and dusk/dawn *Anopheles darling* i.

|  | Indoor<br>Dawn | Indoor<br>Dusk | Outdoor<br>Dawn |
| --- | --- | --- | --- |
| Indoor Dusk | 0.155 |  |  |
| Outdoor Dawn | 0.134 | 0.102 |  |
| Outdoor Dusk | 0.259 | 0.139 | 0.081 |

## Discussion

Reduced representation genome-sequencing methods, such as ddRAD, have proved powerful to assess a large number of SNPs on a genome-wide scale, with considerable lower library construction and sequencing effort. However, these approaches suffer from sampling biases i.e., allele dropout (ADO), due to the absence or polymorphism on a restriction site [35]. *In silico* simulations showed that pruned datasets with incomplete loci sampling i.e., missing data are more likely to accurately reflect population genetic [36]. Here, we applied two strict filters for all set of analysis: no prior information of population and loci were selected when presented in at least 50% of individuals.

In this study, we investigated the genetic basis of biting behavior regarding location (indoor/outdoor) and time (dusk/dawn) in females of *An. darlingi*. There was evidence of a genetic component for both local and time biting behaviors features using genome-wide SNPs. In the Peruvian Amazon, there was no evidence of genetic differentiation between exophagic and endophagic *An. darlingi [29]*. Hence, this is the first study to report a genetic component related to *An. darlingi* behavior. Previously, studies with other species such as *An. gambiae* s.l. and *An. funestus* showed genetic differentiation regarding behavior using chromosomal inversion frequencies [37,38]. In addition, a genetic component was detected for host choice for blood-feeding in *An. arabiensis* using whole-genome sequencing [39]. However, this species showed no evidence of genetic differentiation for biting and resting behavior [39,40].

Concerning location, females collected outdoor in two different rural settlements (Granada and Remansinho), about 60 km apart, comprised two genetic clusters. This finding corroborated prior reported of microgeographic genetic differentiation studies found in *An. darlingi* by habitat [18]. On the other hand, these two independent places presented a similar indoor population of *An. darlingi* by PCA, DAPC and presented the lowest pairwise *F_ST_* (Fig 2). This single indoor group could be explained by similar selection of *An*. *darlingi* female related to environmental conditions within houses such as temperature, sheltering and blood meal availability. In fact, Paaijmans and Thomas [41] showed that indoor environments are warmer than outdoor ones and usually presented less daily variation. Besides temperature, indoor residual spraying could be an important drive in the selection of mosquito females. Different reports already showed fast selection of mosquito resistance genes correlated to insecticide usage, either by insecticide-treated bed nets (ITNs) and indoors residual spraying [42–45]. Both rural settlement studied (Granada and Remansinho) have no long history in use of ITNs, but both places usually use indoors residual spraying.

In the matter of the time of biting behavior in *An. darlingi,* the present study showed non-random distribution of individuals in the PCA and clustering in DAPC. As shown in the first set of analysis, PC1 segregated indoors and outdoors individuals (Fig 3). In addition, PC2 segregated dusk and dawn individuals. In fact, DAPC presented four genetic clusters containing samples from each category. Across its broad distribution, *An. darlingi* populations present a range of peak biting times and patterns i.e., unimodal, bimodal, trimodal [21,24]. Biting time variation of *An. darlingi* showed associated with seasonality [19,46] and local and ecological factor [47]. The current study collected *An. darlingi* biting throughout the night with two peaks, one in the dusk and another in the dawn (data not published, supplementary). Prussing and collaborators [29] found no evidence of genetic differentiation between *An. darlingi* biting times during the night in Iquitos, Peru. However, *An. darlingi* have presented especially early-evening biting activity in that area [48,49], which disagreed with the pattern found in the present study. Additional sampling could help to determine whether the differentiation detected in this study is consistent between populations of *An. darlingi* showing similar behavior patterns.

It is widely accepted that behavioral aspects of anopheline species such as host preference, biting time and resting location after a blood meal have a major impact on malaria transmission dynamics in endemic areas [50,51]. From an evolutionary point of view, it is tempting to assume that during *An. darlingi* colonization of the Amazon region, mosquito subpopulations began to evolve in order to occupy the myriad niches in this diverse environment. Together with recent events directly attributable to human presence such as deforestation and land use changes on a massive scale [9,21], we hypothesize that these processes have led to structured populations of *An. darlingi* that may at least partially explain the strong evidence for genetic and behavioral diversity. Besides deflorestation, another important human modification in Amazon region is the construction of small pods to supplement water all over the year. This permanent water collection near the houses could be an important breeding site to support anophelines populations with close access to human host. A better understanding of the genetic basis for diversity in *An. darlingi* may help to predict and improve vector control methods and effectiveness of malaria frontline interventions.

## Materials & Methods

### Samples

Collections of *Anopheles darlingi* adults were performed outdoors (peridomestic, within 10m of each house) and indoors in two rural settlements, Granada (9°45’S 67°04’W) and Remansinho (9°29’S 66°34’W) in Acre State in April 2011(Fig 1). Collections were performed using human landing catch (HLC) by the authors MC and PEMR, between 18-06 h. All collected specimens were identified using the key of Consoli & Lourenço-Oliveira [30] based on morphological characters and conserved individually in eppendorf tubes at -20ºC; only *An. darlingi* specimens were used for further analysis.

### SNP genotyping

DNA was individually extracted using ReliaPrep™ Blood gDNA kit (Promega, Madison, USA) and DNA concentration was estimated using a Qubit fluorometer (Invitrogen, Carlsbad, USA). Double restriction digestion of 200 ng of high quality genomic DNA with *EcoR*l-*Msp*l restriction enzymes was performed in a 40 ul reaction volume and then purified with AMPure XP beads following the manufacturer’s protocol (Beckman Coulter, California, USA). Hybridized customized adapters P1 (0.3uM) and P2 (4.8uM) were ligated to the digested DNA (T4 DNA Ligase, Promega) as described in Campos et al. (2017). After another purification with AMPure XP beads, DNA was size selected on an agarose gel to 350-400 bp followed by another AMPure XP bead purification. PCR amplification for Nextera^®^ indexing was carried out to generate Illumina sequencing libraries, according to these cycling conditions: an initial denaturation step at 72ºC for 3 min and at 95ºC for 30s, followed by 16 cycles of 95ºC for 10s, annealing at 55ºC for 30s, elongation at 72ºC for 30s, and a final extension cycle at 72ºC for 5 minutes, then each PCR product was purified one last time. Samples were individually quantified using the KAPA library quantification kit (KAPA Biosystems, Wilmington, USA) and equimolarly combined to compose a final library. Final libraries were quantified, normalized, denatured, and finally loaded on the Illumina reagent cartridge 150-cycle of paired-end sequencing in a Hiseq 2500 (Genomics and Bioinformatics Core, State University of New York at Buffalo).

### SNP data processing

Raw ddRAD reads were processed to identify SNP loci within and between individuals using scripts implemented in Stacks v1.31 [31]. First, sequences were quality filtered using the default parameters of *process_radtags* script. Then, each individual’s sequence reads were aligned to the *An. darlingi* reference genome [32] using Bowtie2 with default parameters [33]. Aligned reads were input to *ref_map.pl* Perl script in Stacks, using a minimum of 5 reads (*-m*) to report a stack and join the catalog. The dataset was corrected using another Stacks script called *rxstacks* with the following parameters: prune out non-biological haplotypes unlikely to occur in the population (*--prune_haplo*), minimum log likelihood required was -10.0 (*--lnl_lim*). A new catalog was built by *cstacks* Stacks’ script and each individual was matched against the catalog with *sstacks* script. We then used the *populations* script in Stacks to filter loci that were called in at least 50% (*-r*) of all samples i.e., the first step was performed without population information to avoid population bias in the SNP selection. This latter step was run with minimum coverage of 5 (-m) and a random single SNP for each RAD locus was selected (*write_random_snp*). The *populations* script produced genotype output in several formats (e.g. VCF, Genepop) and summary statistics such as nucleotide diversity, pairwise *F_ST_* and private alleles.

### SNP data analysis

Principal component analysis (PCA) and discriminant analysis of principal components (DAPC) were performed in the R package Adegenet [34]. The former described global diversity overlooking group information whereas DAPC maximize differences between groups and minimizes variation within clusters. The optimum number of genetic clusters in DAPC was the lowest Bayesian Information Criterion (BIC) associated with several *K*-means calculated.

### Abbreviations

BIC: Bayesian information criterion
DAPC: Discriminant analysis of principal components
ddRADseq: Double digestion restriction-site associated DNA sequencing
PCA: Principal component analysis
SNP: single nucleotide polymorphism

### Author contributions

PEMR and MC designed the field and laboratory work; MC and DPA performed the laboratory research; PEMR, MC and KJE analyzed data. All authors actively contributed to the interpretation of the findings; PEMR, MC, DPA wrote, JEC and KJE revised the manuscript. All authors approved the final manuscript.

### Funding

MC was supported by FAPESP. JMV received funding from International Centers for Excellence in Malaria Research  (grant U19AI089681). KJE received funding St. Mary’s College of Maryland. PEMR has a CNPq fellowship.

## Supporting Information Legends

S1 Table. Per-individual *An. darlingi* detail of the number of sequence reads and unique stacks genotyped.

